# Genome-wide SNP genotyping as a simple and practical tool to accelerate the development of inbred lines in outbred tree species: an example in cacao (*Theobroma cacao* L.)

**DOI:** 10.1101/2022.06.12.495796

**Authors:** Uilson Vanderlei Lopes, Jose Luis Pires, Karina Peres Gramacho, Dario Grattapaglia

**Affiliations:** Cacao Research Center (CEPEC/CEPLAC), Rod. Ilhéus-Itabuna, km 22, Ilhéus, BA, Brazil; Plant Genetics Laboratory, EMBRAPA Genetic Resources and Biotechnology, CEP 70770-970, DF, Brasilia, Brazil

**Keywords:** *Theobroma cacao*, SNP, breeding, hybrid, inbred lines, homozygosity

## Abstract

Cacao is a globally important crop with a long history of domestication and selective breeding. Despite the increased use of elite clones by cacao farmers, worldwide plantations are established mainly using hybrid progeny material derived from heterozygous parents, therefore displaying high tree-to-tree variability. The deliberate development of hybrids from advanced inbred lines produced by successive generations of self-pollination has not yet been fully considered in cacao breeding. This is largely due to the self-incompatibility of the species, the long generation cycles (3-5 years) and the extensive trial areas needed to accomplish the endeavor. We propose a simple and accessible approach to develop inbred lines based on accelerating the buildup of homozygosity based on regular selfing assisted by genome-wide SNP genotyping. In this study we genotyped 90 clones from the Brazilian CEPEC’s germplasm collection and 49 inbred offspring of six S_1_ or S_2_ cacao families derived from self-pollinating clones CCN-51, PS-13.19, TSH-1188 and SIAL-169. A set of 3,380 SNPs distributed across the cacao genome were interrogated on the EMBRAPA multi-species 65k Infinium chip. The 90 cacao clones showed considerable variation in genome-wide SNP homozygosity (mean 0.727± 0.182) and 19 of them with homozygosity >90%. By assessing the increase in homozygosity across two generations of self-pollinations, SNP data revealed the wide variability in homozygosity within and between S_1_ and S_2_ families. Even in small families (<10 sibs), individuals were identified with up to ∼1.5 standard deviations above the family mean homozygosity. From baseline homozygosities of 0.476 and 0.454, offspring with homozygosities of 0.862 and 0.879 were recovered for clones TSH-1188 and CCN-51 respectively, in only two generations of selfing (81-93% increase). SNP marker assisted monitoring and selection of inbred individuals can be a practical tool to optimize and accelerate the development of inbred lines of outbred species. This approach will allow a faster and more accurate exploitation of hybrid strategies in cacao breeding programs and potentially in other perennial fruit and forest trees.

## Introduction

Cacao (*Theobroma cacao* L.) is a predominantly allogamous tropical tree species [1], whose beans are the major ingredient for the chocolate industry. Since the report of heterosis in cacao [2–4], the crop is planted worldwide mainly as full-sib families of interclonal hybrids, although clones are also widely used in some countries [5]. Clones used as parents in the production of cacao hybrid progenies are frequently heterozygous and, as a consequence, the plantations suffer of an undesired tree-to-tree variability [6]. Alternatives to reduce such variability have been proposed through the use of clones or hybrids between partially or fully inbred lines.

Cacao clones are largely used in countries like Brazil (especially Bahia and Espírito Santo states), Ecuador, Malaysia, Indonesia, among others. However, despite the many benefits of cloning, there are also some drawbacks [7]. First, the process of grafting and managing clones is not easy, particularly for unskilled small farmers. Second, most propagules (budsticks and cuttings) used in cacao cloning comes from plagiotropic branches, resulting in plants with an architecture that requires intensive pruning, especially for clones with a prostrate habit (ex. Scavinas). Third, plagiotropic clones usually do not have a true tap root, potentially precluding adequate establishment in areas suboptimal for cacao. Fourth, delivering propagation material (budsticks) is not easy compared to seeds, particularly in remote areas. In order to minimize some of those problems, somatic embryo plants have been suggested [8], but besides the genotype dependence for the success of this method, the lab facilities required to produce somatic plants are beyond the budget of most producer countries. Today only Indonesia has planted somatic cacao plants in a moderately large scale. Other more promising strategies for third world countries (e.g., combining tissue culture and increased production of orthotropic propagules) have also been proposed [8] and can facilitate its adoption. But some of the challenges still persist.

Hybrid cacao varieties are planted in most producer countries in the world, including those in West Africa, where around 70% of the world cocoa beans are produced [9]. Besides the potential exploration of heterosis, hybrids offer additional advantages when compared to clones. Not only are they planted by seeds, facilitating propagation by farmers, but they also improve establishment in the field and eventually water absorption in deep regions of the soil due to their tap roots. Additionally, hybrid plants benefit from an orthotropic architecture that facilitates management (e.g., pruning, movement in the area), especially for farmers already used to the orthotropic habit of local open-pollinated varieties. On the other hand, because cacao hybrids are produced mostly by crossing heterozygous clones, the trees display an unwanted variability in yield and vigor [6]. The differences in vigor among trees increase competition among them, resulting in lower yield and eventually death of the weaker plants and a reduced overall stand after some years.

The use of inbred lines as parents of hybrids could minimize the problems associated with both, clonal propagation and interclonal hybrid variability, by consolidating the advantages of both deployment methods, i.e., seed propagation and capture of hybrid vigor. This was attempted since the beginning of hybrid breeding in cacao by searching for naturally inbred selections from local populations (e.g., Amelonado, Matina) [10–13], using partially inbred lines [11, 13–15], looking for spontaneous haploids to be diploidized [16] and through the use of anther culture [17–19]. All these strategies ultimately try to overcome the challenge associated with the production of fully inbred parents in cacao by the traditional method of successive self-fertilizations, a lengthy process, given the long generation time in cacao (3-5 years/generation) and large experimental area required (commonly 9 m^2^/tree).

Genome-wide DNA marker data provides a simple and accurate proxy of the genome-wide level of homozygosity of an individual plant produced by self-pollination by simply counting the proportion of marker loci that, once heterozygous in the self-pollinated parent, segregated to homozygosity in the inbred offspring. Individuals with higher levels of attained homozygosity would be prioritized to carry out the subsequent generation of self-pollination, fast-tracking the production of inbred lines when compared to simple random selection of offsprings. Despite this obvious application of DNA marker data to accelerate the buildup of inbreeding in outbred plants, reports to this end have been rare, using only low-resolution microsatellite genotyping in papaya [20] and cassava [21]. In both studies, plants with higher-than-expected homozygosity could be identified already in the initial generations of self-pollination.

With the dramatic reduction of sequencing and genotyping costs in recent years, cacao has experienced significant advances in genomic resources and knowledge [9]. Two cacao genomes have been sequenced and assembled [22, 23], thousands of SNPs were discovered and gathered in user-friendly, high-precision genotyping platforms [24–26], and a large number of germplasm accessions genotyped. Despite the many applications of SNPs in cacao genetics and breeding, including diversity studies [27–30], QTL mapping, genome-wide association and genomic selection [31–36], they have not been used to accelerate the development of inbred lines. Nevertheless, population and individual level estimated heterozygosity have been published for cacao germplasm collections and wild populations using microsatellites [37–40] or hundreds of SNPs [41, 42]. Large scale SNP heterozygosity surveys have been published for a few key clones from transcriptome data [26] or full genome sequencing [43]. Variable levels of heterozygosity were found across single clones and populations, with some of them displaying significant heterozygosity deficiency. This same data, when looked from the alternative perspective of inbreeding, have therefore revealed variable levels of homozygosity in the existing germplasm, with some highly homozygous clones.

In this study we show that genome-wide SNP marker data can be efficiently used to assess the increase in homozygosity across two generations of self-fertilizations, revealing a wide variability in the attained homozygosity among S_1_ and S_2_ individuals within and across families. Furthermore, the genome-wide levels of homozygosity in a set of germplasm accessions of the Cacao Research Center (CEPEC/CEPLAC) in Brazil were estimated to potentially select which clones to prioritize for the development of inbred lines, as well as to drive pairwise clone combinations to generate hybrid progenies with higher levels of heterozygosity and potentially heterosis.

## Material and methods

### Germplasm accessions and inbred progenies

A diverse sample of 90 cacao clones was selected to estimate the levels of individual homozygosity at SNP markers. The accessions are part of CEPEC’s germplasm collection, which currently hosts around 2,000 entries. The 90 clones were chosen to represent the subpopulations described earlier [44] and/or because of their higher relevance to CEPEC’s breeding program. Among those are wild germplasm and breeder or on-farm selections. Several of the 90 clones sampled in this study have been used to establish a program aiming at the development of advanced inbred lines of cacao for future production of hybrid seeds. In that program 31 S_1_ families were produced with 16 to 53 individuals per family (average= 32.8 trees/family), and eight S_2_ families, with 26 to 33 individuals per family (average=30.1 trees/family). The S_1_ and S_2_ families were generated by selfing S_o_ and S_1_ plants, respectively, by manual (protected) pollination, as usually done in cacao. Among those 90 clones, four are widely used in CEPEC’s breeding program, because of their high yield and/or resistance to diseases: CCN-51, PS-13.19, TSH-1188 and SIAL-169. Six inbred families were genotyped in this study to evaluate the feasibility of the proposed approach of assessing the within-family variability in homozygosity and identify more homozygous offspring. Two S_1_ families were obtained by selfing cacao clones PS-13.19 and SIAL-169; four S_2_ families, three of them obtained by selfing three different S_1_ offsprings of clone TSH-1188; and one family from an S_1_ offspring of clone CCN-51 (Fig 1).

**Fig 1.**
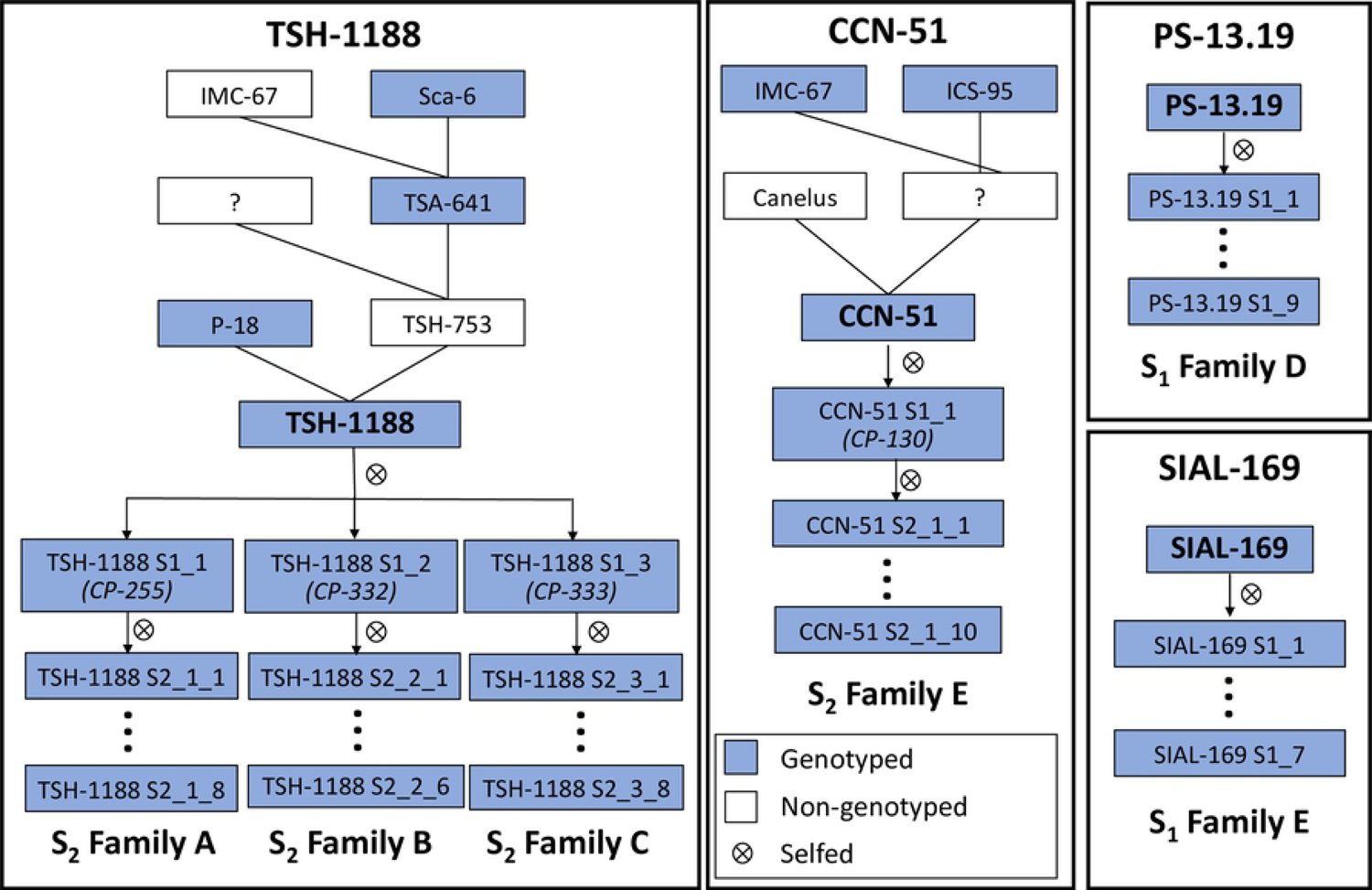
Pedigrees of the six partially inbred families obtained from self-pollination of four cacao clones (TSH-1188, CCN-51, PS-13.19, SIAL-169).

The seeds produced were planted in 288 cm^3^ containers with fertilized soil. Around 5 months after seeding, the plants were established in a progeny trial together with progenies involving sources of resistance to moniliasis (*Moniliophthora roreri*) (trial PT-1501). PT-1501 was established using a spacing of 3.0 x 1.5 m, in June/2015, under a randomized complete block design, with 3 blocks and 21 plants per plot, and 102 progenies involving sources of resistance to moniliasis (6,804 plants). In some plots of that trial, the 1,261 S_1_ and S_2_ individuals were planted. To assess the prospects of the approach proposed in this study, between 6 to 9 inbred plants per family showing good performance were genotyped (Fig 1).

### SNP genotyping

Mature and healthy leaf samples were collected from each one of the S_1_ and S_2_ offspring individuals of the inbred families in trial TP-1501 together with the 90 clones from CEPEC’s germplasm collection in Ilhéus, BA. Leaf samples were stored in 50 ml plastic tubes with silica gel until use. Total genomic DNA was extracted with an optimized protocol that uses a pre-wash with a sorbitol buffer [45] and quantified with a Nanodrop 2000 spectrophotometer (Thermo Fisher Scientific, MA). A set of 3,412 SNPs were selected from those validated in the previously developed 6K and 15K “chocolate” SNP chips (Tcm SNPs) [25, 26], to populate a sector of the EMBRAPA multispecies 65K Infinium chip. This chip contains a total of 66,413 SNPs, shared among 27 different plant and animal species, significantly reducing the individual sample genotyping cost, while at the same time allowing the generation of high-quality and inter-laboratory portable SNP data for all species (Grattapaglia D. et al. in preparation). Cacao SNPs were selected based on a set of criteria that included performance metrics of the SNPs in previously genotyped germplasm, including SNP call rate (CR), Minimum allele frequency (MAF), SNP quality parameters (e.g. GenomeStudio GenCall score) from previous reports [25, 26]. Information regarding SNP genome address in the reference genome was also taken into account to distribute SNPs across chromosomes to cover as much as possible the recombination space and allow genotype imputation in upcoming studies. To optimize the available space on the chip, only Infinium II SNPs were chosen, therefore genotyping only four out of the six possible SNPs configurations, namely, A/G, A/C, C/T and G/T. Final evaluation of the selected SNP probeset included a detailed sequence evaluation to avoid SNP probe cross-talking with the genome of the other species. Ultimately a selected set of 3,412 cacao SNP probes were placed on the multispecies chip (S1 File).

Genotyping was carried out at Neogen/Geneseek (Lincoln, NE). Manifest files and intensity data (.idat files) were obtained from Neogen. SNP genotypes were called using GenomeStudio 2.0 (Illumina, Inc. San Diego, CA) following the standard genotyping and quality control procedures [46] and exported in the AB format where alleles A and T at the SNPs are coded as “A” and alleles G or C at the SNPs are coded as allele “B”. As a quality control measure for the SNP data, five cacao clones (CCN-51, CSUL-3, RB-45, SCA-6 and TSH-1188) were genotyped with replicate samples.

### Data analysis

A parentage test using the SNP marker data was carried out on all S_1_ or S_2_ individuals prior to subsequent analyses. Given the focus of the study (inbred line development), all analyses were done in terms of the observed and expected homozygosities, and not in terms of heterozygosities as usually done in diversity studies. For each individual plant the number of homozygous (AA and BB) and heterozygous (AB) SNPs genotypes were counted and the observed homozygosity estimated as the total proportion of homozygous SNPs over the total number of genotyped SNPs. For each inbred S_1_ or S_2_ family, the mean (μ) and standard deviation (σ) of the observed homozygosity were calculated. Individual offspring homozygosities were normalized by the standard deviation (σ) to allow ranking and identifying the S_1_ and S_2_ individuals with higher proportions of homozygosity within each family. A chi-square test (α = 0.05) was used to test the null hypothesis of no difference between the observed and expected counts of the three SNP genotypic classes (AA, AB, BB) for each offspring individual. Expected genotypic counts for the inbred generation were obtained based on a simple Mendelian model assuming no selection. In other words, from the genotypic counts (AA, AB, BB) in the self-pollinated parent, the number of homozygous SNPs (AA and BB) counts are expected to increase by 25% each, and the number of heterozygous SNP counts to decrease by 50% in each generation. A significant chi-square would indicate either a less than expected or higher than expected homozygosity from Mendelian expectations, providing a statistical assessment of the deviation due to sampling effects or selection against inbreeding in each individual offspring.

The average expected homozygosity in each inbred family was estimated in an analogous manner, i.e., based on the observed homozygosity of the self-pollinated parent. In other words, the homozygosity is expected to increase by 50% of what was the observed homozygosity in the previous generation. For one of the S_2_ families (family C) the S_1_1_ parent (CP-255, Fig 1) was missing. The average observed homozygosity of its siblings S_1_2_ and S_1_3_ was used instead for the calculations. A t-test of the null hypothesis of no difference between the family mean observed and expected numbers of homozygous SNP counts was used to assess the deviation from the expected overall level of inbreeding in the family. Finally, the root-mean-square-deviation (RMSD) between the observed and Mendelian model predicted homozygosity was also estimated for each family, as a way to quantify the deviation from the predicted inbreeding.

A matrix of genetic distances based on the SNP data among all 90 clones and the S_1_ and S_2_ individuals was calculated as a preliminary tool to choose what crosses among them would yield hybrids with the highest proportions of SNP loci in heterozygosity. Genetic distances were estimated using the genetic distance measure of Smouse and Peakall [47] between a pair of individuals for a codominant locus using Genalex 6.5 [48]. UPGMA dendrograms based on the genetic distance matrix for the 90 clones alone, and the 90 clones plus the S_1_ and S_2_ individuals were built using MegaX [49] to visualize the genetic relationships among them.

## Results

### SNP data

Raw SNP data exported from GenomeStudio 2.0 with GenCall>0.15 were re-clustered and submitted to additional quality controls. Only SNPs that passed the following Illumina multi-variable metrics criteria were retained: (i) genotype clusters separation > 0.3; (ii) mean normalized intensity (R) value of the heterozygote cluster >0.2; (iii) mean normalized theta of the heterozygote cluster between 0.2 and 0.8; (iv) >99% reproducibility across the replicated samples and >99% correct inheritance between generations in the inbred families. Only SNPs with call rate ≥95% and MAF≥ 0.01 for samples with call rate ≥87% were retained for further analyses. Out of the 3,412 SNPs present on the chip, data for 3,380 SNPs were ultimately retained after quality control. The full data set is provided (S2 File). Although a MAF cutoff of 0.01 was used, the vast majority of the SNPs (3,341 of the 3,380, 98.8%) had MAF > 0.05 and the whole site frequency spectrum showed a negatively left-skewed distribution toward higher MAF values (mean = 0.325±0.123; median= 0.347; S3 Fig) with 2,442 SNPs (72.2%) with MAF ≥0.25 estimated using only the set of 90 clones.

### Cacao clones

The average observed homozygosity for this group of 90 clones was 0.727±0.182 (S4 File) while the expected homozygosity under Hardy Weinberg equilibrium was 0.591. A total of 2,771 SNPs deviated from HWE expectations at the nominal p-value ≤ 0.05 and 477 SNPs at the Bonferroni corrected p-value ≤ 1.47E-05 for multiple tests. A highly significant inbreeding coefficient (F= 0.323 ±0.002; p-value<0.0001) was therefore estimated. The 90 cacao clones showed considerable variation in genome-wide SNP homozygosity, although negatively left-skewed toward higher values (mean = 0.727; median= 0.768; Fig 2, S4 File). Nineteen clones showed homozygosities above 90% and 10 of them above 95%: APA-4, SIC-806, CAB-44, SIC-2, GU-123C, SIAL-169, CAIRU-1, CAIRUIGREJA-1, NA-286 and SIC-20. Some clones, widely used in many breeding programs worldwide, including SCA-6, ICS-1, ICS-6, P-7, NA-33 and IMC-67, showed homozygosity estimates ranging from 0.558 to 0.845. The lowest estimates of homozygosity (≤ 0.40) were found for clones CC-10, ICS-95, ICS-7. Among the 90 clones, a set of 62 wild/semi-wild accessions including some Amelonado selections from old plantations (SIC and SIAL series clones) displayed homozygosities above 0.61, averaging 0.734 ± 0.175, contributing to a large extent for the skewed distribution toward higher homozygosity values (Fig 2).

**Fig 2.**
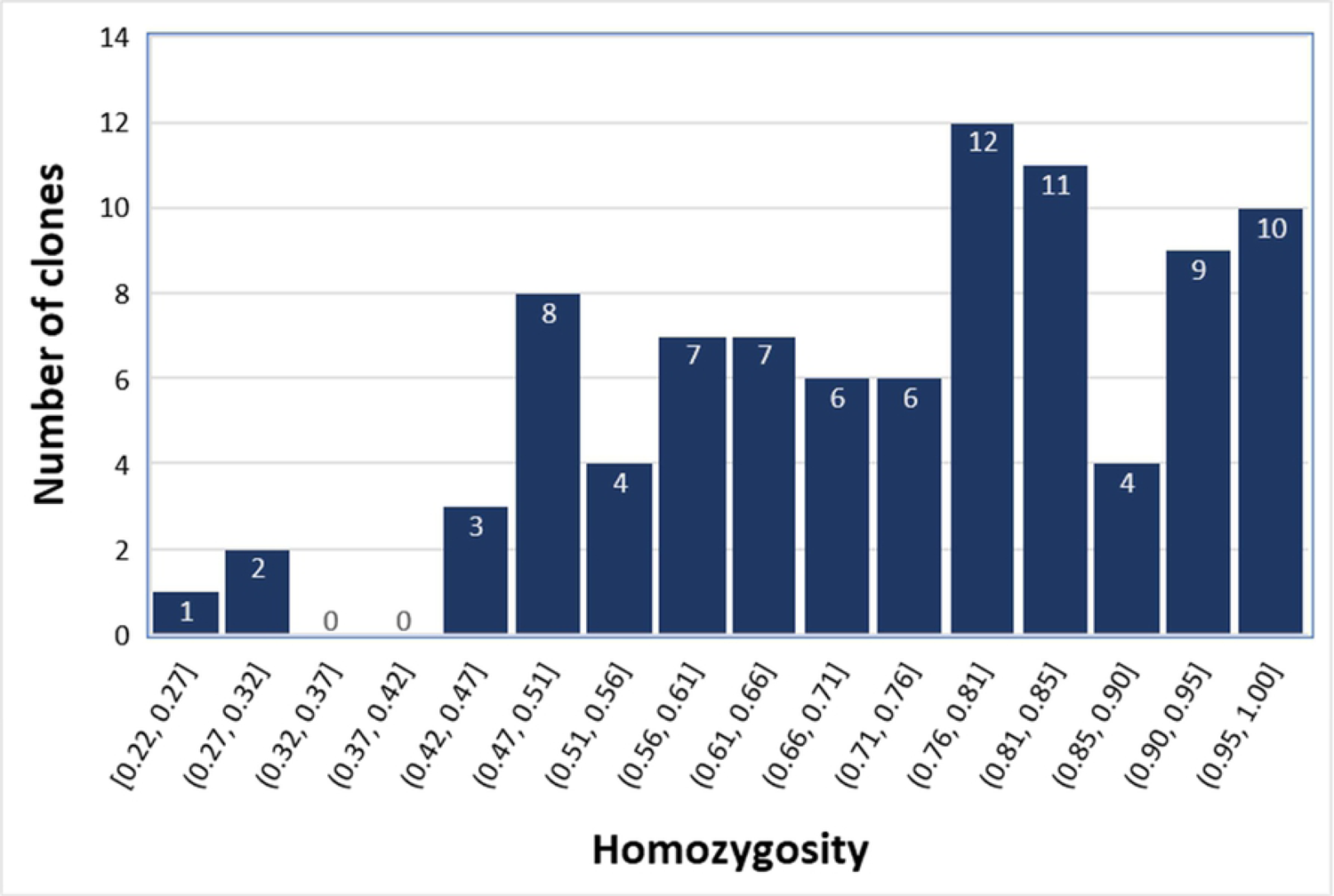
Distribution of the genome-wide homozygosity of the 90 cacao clones at the 3,380 sampled SNP

A matrix of genetic distance analysis was built providing estimates of genome-wide genetic distance among the 90 clones together with the S_1_ and S_2_ inbred individuals studied (see below) (S5 File). UPGMA dendrograms based on the genetic distance matrix were built for the 90 clones only, and the 90 clones plus the 49 S_1_ and S_2_ individuals. The 90 clones were positioned into clusters for the most part consistent with expectations, although three clones clearly did not fit the expected groupings, most likely reflecting mislabeling problems. These were clones SCA-12, SIAL-70 and CCN-16 (S6 Fig).

### S_1_ and S_2_ inbred families

Upon parentage testing, three S_2_ individuals supposedly derived from clone TSH-1188 were excluded from the study based on the observation of several hundred SNP genotypes incompatible with parent-offspring inheritance. For all remaining 49 confirmed inbred plants, 31 S_2_ and 18 S_1_ plants, estimates of individual (Table 1) and family (Table 2) SNP homozygosities were calculated. Except for parent SIAL-169 that showed an already high homozygosity of 0.969, the starting homozygosity in the other three S_0_ parental clones (TSH-1188, PS-13.19 and CCN-51) was around 0.5 ranging from 0.454 for CCN-51 to 0.491 for PS-13.19. Given the very high homozygosity of clone SIAL-169, its S_1_ inbred family showed little variation in homozygosity with the average, minimum and maximum values close to the homozygosity of the original SIAL-169 parent. As expected, for the other five families, the range and the average homozygosity in the four S_2_ families was higher than the homozygosity seen in the S_1_ family. This S_2_ to S_1_ difference in homozygosity was smaller for the three S_2_ families of clone TSH-1188 and larger for the S_2_ family of clone CCN-51. While the average homozygosity was 0.867 in the S_2_ family of CCN-51, and between 0.753 and 0.786 in the three S_2_ families of clone TSH-1188, it was 0.715 in the S_1_ family of clone PS-13.19. The higher average homozygosity of the S_2_ family of CCN-51 (0.867) is explained by the higher homozygosity (0.741) already seen in the S_1_ plant (CCN-51_S1_1) when compared to the S_1_ plants of clone TSH-1188. Overall, no significant difference was seen between the observed and expected family mean homozygosity (all t-tests non-significant) and the deviations (RMSD) between the Mendelian model predicted and the observed homozygosity were small, varying from 0.3% for the S_1_ family of SIAL-169, to a maximum of 6% for the S_1_ family of PS-13.19 (Table 2).

**Table 1.** Summary of the SNP genotyping data for the individual offsprings in the six inbred families produced by self-fertilization of four cacao clones indicated as S_0_ parents (TSH-1188, PS-13.19, CCN-51 and SIAL-169). Reported are the genotypic counts, the observed and expected homozygosities, standardized homozygosity and the chi-square test for departure from the average homozygosity by Mendelian expectation.

| Inbred Family | Generation | Individual | # SNP AA | # SNP AB | # SNP BB | Total # | Obs. Hom. | Exp. Hom. | Std. Hom. | chi-square |
| --- | --- | --- | --- | --- | --- | --- | --- | --- | --- | --- |
| A | S0 | TSH-1188 | 812 | 1770 | 798 | 3380 | 0.476 | - | - | - |
| A | S1 | TSH-1188_S1_2 | 924 | 1542 | 914 | 3380 | 0.544 | 0.738 | - | 660.74** |
| A | S2 | TSH-1188_S2_2_2 | 1193 | 1044 | 1141 | 3378 | 0.691 | 0.772 | -1.879 | 126.36** |
| A | S2 | TSH-1188_S2_2_1 | 1285 | 817 | 1267 | 3369 | 0.757 | 0.772 | -0.558 | 4.02ns |
| A | S2 | TSH-1188_S2_2_3 | 1317 | 746 | 1317 | 3380 | 0.779 | 0.772 | -0.126 | 1.09ns |
| A | S2 | TSH-1188_S2_2_7 | 1295 | 740 | 1343 | 3378 | 0.781 | 0.772 | -0.093 | 2.86ns |
| A | S2 | TSH-1188_S2_2_6 | 1285 | 729 | 1363 | 3377 | 0.784 | 0.772 | -0.030 | 5.85ns |
| A | S2 | TSH-1188_S2_2_5 | 1350 | 670 | 1359 | 3379 | 0.802 | 0.772 | 0.319 | 17.21** |
| A | S2 | TSH-1188_S2_2_8 | 1387 | 580 | 1409 | 3376 | 0.828 | 0.772 | 0.844 | 61.13** |
| A | S2 | TSH-1188_S2_2_4 | 1474 | 465 | 1440 | 3379 | 0.862 | 0.772 | 1.523 | 157.3** |
| B | S0 | TSH-1188 | 812 | 1770 | 798 | 3380 | 0.476 | - | - | - |
| **B** | **S1** | **TSH-1188_S1_3** | 909 | 1536 | 934 | 3379 | 0.545 | 0.738 | - | 649.75** |
| **B** | **S2** | **TSH-1188_S2_3_3** | 1186 | 998 | 1193 | 3377 | 0.704 | 0.773 | -1.059 | 89.59** |
| **B** | **S2** | **TSH-1188_S2_3_4** | 1183 | 958 | 1238 | 3379 | 0.716 | 0.773 | -0.797 | 61.22** |
| **B** | **S2** | **TSH-1188_S2_3_1** | 1219 | 954 | 1206 | 3379 | 0.718 | 0.773 | -0.771 | 58.8** |
| **B** | **S2** | **TSH-1188_S2_3_5** | 1314 | 752 | 1311 | 3377 | 0.777 | 0.773 | 0.529 | 0.71ns |
| **B** | **S2** | **TSH-1188_S2_3_2** | 1321 | 716 | 1339 | 3376 | 0.788 | 0.773 | 0.760 | 4.46ns |
| **B** | **S2** | **TSH-1188_S2_3_6** | 1386 | 627 | 1366 | 3379 | 0.814 | 0.773 | 1.338 | 34.32** |
| **C** | **S0** | **TSH-1188** | 812 | 1770 | 798 | 3380 | 0.476 | - | - | - |
| **C** | **S1** | **TSH-1188_S1_1\*** | - | - | - | - | 0.545 | 0.738 | - |  |
| **C** | **S2** | **TSH-1188_S2_1_1** | 1250 | 880 | 1249 | 3379 | 0.740 | 0.773 | -1.620 | 21.38** |
| **C** | **S2** | **TSH-1188_S2_1_7** | 1269 | 827 | 1282 | 3378 | 0.755 | 0.773 | -1.103 | 5.96ns |
| **C** | **S2** | **TSH-1188_S2_1_8** | 1335 | 702 | 1332 | 3369 | 0.792 | 0.773 | 0.103 | 7.18** |
| **C** | **S2** | **TSH-1188_S2_1_6** | 1345 | 691 | 1344 | 3380 | 0.796 | 0.773 | 0.233 | 10.32** |
| **C** | **S2** | **TSH-1188_S2_1_5** | 1335 | 665 | 1352 | 3352 | 0.802 | 0.773 | 0.433 | 16.06** |
| **C** | **S2** | **TSH-1188_S2_1_4** | 1396 | 615 | 1368 | 3379 | 0.818 | 0.773 | 0.975 | 40.58** |
| C | S2 | TSH-1188_S2_1_3 | 1316 | 589 | 1333 | 3238 | 0.818 | 0.773 | 0.979 | 42.3** |
| D | S0 | PS-13.19 | 813 | 1721 | 846 | 3380 | 0.491 |  | - |  |
| D | S1 | PS-13.19_S1_1 | 1051 | 1266 | 1063 | 3380 | 0.625 | 0.745 | -1.622 | 256.45** |
| D | S1 | PS-13.19_S1_5 | 1058 | 1184 | 1138 | 3380 | 0.650 | 0.745 | -1.182 | 164.2** |
| D | S1 | PS-13.19_S1_4 | 1123 | 1093 | 1158 | 3374 | 0.676 | 0.745 | -0.705 | 85.41** |
| D | S1 | PS-13.19_S1_7 | 1150 | 966 | 1226 | 3342 | 0.711 | 0.745 | -0.073 | 21.91** |
| D | S1 | PS-13.19_S1_6 | 1192 | 934 | 1252 | 3378 | 0.724 | 0.745 | 0.155 | 8.85** |
| D | S1 | PS-13.19_S1_3 | 1215 | 888 | 1275 | 3378 | 0.737 | 0.745 | 0.401 | 1.52ns |
| D | S1 | PS-13.19_S1_2 | 1211 | 829 | 1340 | 3380 | 0.755 | 0.745 | 0.720 | 5.17ns |
| D | S1 | PS-13.19_S1_8 | 1262 | 791 | 1324 | 3377 | 0.766 | 0.745 | 0.920 | 7.68** |
| D | S1 | PS-13.19_S1_9 | 751 | 619 | 1598 | 2968 | 0.791 | 0.745 | 1.385 | 343.79** |
| E | S0 | CCN-51 | 767 | 1846 | 767 | 3380 | 0.454 | - | - |  |
| E | S1 | CCN-51_S1_1 | 1239 | 874 | 1267 | 3380 | 0.741 | 0.727 | - | 3.9ns |
| E | S2 | CCN-51_S2_1_8 | 1415 | 510 | 1455 | 3380 | 0.849 | 0.871 | -1.793 | 14.06** |
| E | S2 | CCN-51_S2_1_5 | 1438 | 507 | 1434 | 3379 | 0.850 | 0.871 | -1.712 | 13.26** |
| E | S2 | CCN-51_S2_1_4 | 1443 | 445 | 1492 | 3380 | 0.868 | 0.871 | 0.074 | 0.32ns |
| E | S2 | CCN-51_S2_1_3 | 1470 | 445 | 1465 | 3380 | 0.868 | 0.871 | 0.074 | 0.54ns |
| E | S2 | CCN-51_S2_1_9 | 1452 | 444 | 1484 | 3380 | 0.869 | 0.871 | 0.102 | 0.13ns |
| E | S2 | CCN-51_S2_1_6 | 1464 | 440 | 1476 | 3380 | 0.870 | 0.871 | 0.217 | 0.11ns |
| E | S2 | CCN-51_S2_1_2 | 1436 | 439 | 1505 | 3380 | 0.870 | 0.871 | 0.246 | 0.58ns |
| E | S2 | CCN-51_S2_1_1 | 1473 | 425 | 1480 | 3378 | 0.874 | 0.871 | 0.641 | 0.51ns |
| E | S2 | CCN-51_S2_1_10 | 1483 | 411 | 1486 | 3380 | 0.878 | 0.871 | 1.050 | 1.99ns |
| E | S2 | CCN-51_S2_1_7 | 1489 | 409 | 1480 | 3378 | 0.879 | 0.871 | 1.101 | 2.5ns |
| F | S0 | SIAL-169 | 1610 | 106 | 1664 | 3380 | 0.969 | - | - |  |
| F | S1 | SIAL-169_S1_1 | 1625 | 72 | 1683 | 3380 | 0.979 | 0.984 | -1.892 | 6.93** |
| F | S1 | SIAL-169_S1_7 | 1638 | 52 | 1690 | 3380 | 0.985 | 0.984 | -0.172 | 0.02ns |
| F | S1 | SIAL-169_S1_6 | 1642 | 49 | 1689 | 3380 | 0.986 | 0.984 | 0.086 | 0.32ns |
| F | S1 | SIAL-169_S1_2 | 1644 | 45 | 1691 | 3380 | 0.987 | 0.984 | 0.430 | 1.24ns |
| F | S1 | SIAL-169_S1_5 | 1641 | 41 | 1697 | 3379 | 0.988 | 0.984 | 0.773 | 2.75ns |
| F | S1 | SIAL-169_S1_3 | 1643 | 41 | 1696 | 3380 | 0.988 | 0.984 | 0.774 | 2.76ns |

**Table 2.** Summary of results of homozygosity of the six inbred cacao families produced by self-fertilization of cacao clones TSH-1188, PS-13.19, CCN-51 and SIAL-169. RMSD: root-mean-square-deviation between the observed and Mendelian model expected homozygosity; t-test of the null hypothesis of no difference between the family mean observed and expected numbers of homozygous SNP counts,

| Inbred family | A | B | C | D | E | F |
| --- | --- | --- | --- | --- | --- | --- |
| Parent | TSH-1188_S1_2 | TSH-1188_S1_3 | TSH-1188_S1_1* | PS-13.19_S0 | CCN-51_S1_1 | SIAL-169_S0 |
| Starting homozygosity of selfed parent | 0.544 | 0.545 | 0.545 | 0.491 | 0.741 | 0.969 |
| # of offspring | 8 | 6 | 8 | 9 | 10 | 6 |
| Type of offspring | S2 | S2 | S2 | S1 | S2 | S1 |
| Expected average homozygosity | 0.772 | 0.773 | 0.773 | 0.745 | 0.871 | 0.984 |
| Observed average homozygosity | 0.786 | 0.753 | 0.789 | 0.715 | 0.868 | 0.985 |
| Min. homozygosity in offspring | 0.691 | 0.704 | 0.740 | 0.625 | 0.849 | 0.979 |
| Max. homozygosity in offspring | 0.862 | 0.814 | 0.818 | 0.791 | 0.879 | 0.988 |
| St.Dev. | 0.050 | 0.046 | 0.030 | 0.055 | 0.010 | 0.003 |
| RMSD | 0.049 | 0.046 | 0.032 | 0.060 | 0.010 | 0.003 |
| t statistics | 0.489 ns | 0.335 ns | 0.373 ns | 0.027 ns | 0.342 ns | 0.572 ns |

\* The S1 parent plant TSH-1188\_S1\_1 was not available. The average genotypic counts of its sibs TSH-1188\_S1\_2 and TSH-1188\_S1\_3 was used as surrogate for the calculation

### S_1_ and S_2_ inbred individuals

There was a wide range in the estimated values of homozygosity across individuals within the S_1_ and S_2_ families, except in the S_1_ family of clone SIAL-169 that was itself already highly homozygous. Individuals with as low as −1.879 and up to +1.523 standard deviations from the mean were recovered (see family A, S_2_ of TSH-1188, Table 1), even with such small numbers of individuals sampled in each family. Individuals within each family were ranked from the lowest to the highest attained homozygosities and a chi-square test on the actual genotypic counts was used to assess the significance of the deviation from the Mendelian expectations for each individual selfed offspring. The magnitudes and significance of the individual chi-square values suggests that the families had different distributions of homozygosity of their offspring (Table 1). For example, for the three S_2_ families of TSH-1188, family A showed a more balanced distribution, with one plant with significantly lower than expected homozygosity, three individuals higher than expected and four individuals within the expected values. On the other hand, family B had four offsprings with lower than expected and only one with higher-than-expected values. Family C had five out of seven individuals with significantly higher than expected homozygosity values. The differences observed could be simply due to a sampling effect, or the result of differential purging of genetic load, or both. Regardless, in all families, the SNP data allowed identifying individual offspring with higher-than-expected homozygosity, although not statistically significant for families D and E. In the proposed approach, these individual plants would be prioritized in the next generations of self-pollination toward an accelerated development of inbred lines.

## Discussion

### Cacao SNP genotyping on the EMBRAPA 65KMultispecies chip

In this study we genotyped a set of 3,412 cacao SNPs selected from the large Illumina validated collections, currently available for cacao [25, 26]. We used an Illumina Infinium fixed-content SNP platform, currently still considered the gold standard in the SNP genotyping industry, despite the rapid growth of alternative sequence-based methods. Following the Illumina recommended quality control thresholds of cluster separation, call rate, genotype reproducibility and inheritance, 3,380 of 3,412 (99%) of the originally interrogated SNPs were retained. Furthermore, SNP quality control of inheritance was also provided by the several parentage verifications done in the inbred families. This result indicates that the SNP performance metrics originally provided by the SNP developers were robust and that our SNP selection for performance efficient. The original choice of cacao SNPs to populate the 65K EMBRAPA multispecies Infinium chip was deliberately made toward highly informative (higher MAF) SNPs, as the main envisaged objectives were genetic mapping, germplasm fingerprinting, parentage testing and characterization. The skewed distribution of the site frequency spectrum observed, toward highly polymorphic SNPs was therefore expected and confirmed (Fig 2). This relatively large set of SNPs should provide a high power of discrimination to carry out a detailed genetic analysis of the entire CEPEC’s cacao collection and breeding populations. Furthermore, it is likely that the 3,412 SNPs used will find overlapping SNPs with other low-density SNP panels used by the main cacao research groups worldwide [24, 27, 50, 51], facilitating data exchange. The fact that these SNPs are part of the large scale EMBRAPA multispecies chip also allowed for a very accessible genotyping cost at around USD 20/sample. The large number of samples assembled across the different plant and animal species encompassed in the EMBRAPA chip, provided the necessary economy of scale for a significant cost reduction on a per sample basis.

### Identity of cacao clones

A considerable number of studies have been published showing a variety of applications of molecular markers, both microsatellites and SNPs in cacao breeding and germplasm characterization (reviewed in [9]). Clone fingerprinting for certification of identity in germplasm collections and progeny trials has been the most operational application. It has revealed that clone mislabeling can be frequent or even pervasive in cacao [38, 40–42, 52]. Mislabeling issues not only seriously affect the efficiency of cacao germplasm conservation and recommendation, but also the correct estimation of genetic parameters in breeding programs [53]. Although fingerprinting was not the main objective of our study, in this relatively limited set of 90 clones surveyed, we detected three clones that did not fit the expected clustering in the UPGMA dendrogram, strongly suggesting a mislabeling problem. Clone SIAL-70 should have clustered in the same groups as the other Amelonado SIC and SIAL clones. Instead, it clustered with clone CCN-16 which is itself also misplaced as it should have clustered close to its parent, clone CCN-51.

Clone SCA-12 was the third case of misplacement. It belongs to the Contamana group and was expected to cluster with its highly related clones SCA-6, SCA-5 and SCA-19. The genetic profile of clone SCA-12 and a few other clones was kindly provided by Dr. Osman Gutierrez (USDA-ARS) as part of a study that established an optimized set of SNPs genotyped by the Agriseq™ genotyping-by sequencing technology [51]. The profile contained 235 Tcm SNPs in common to the 3,380 SNPs genotyped in our study. The comparison revealed 160 mismatching genotypes, confirming the mislabeling of clone SCA-12. Through the same dataset provided by the USDA, it was possible to confirm the matching identity of clones CAB-0224, MA-15, SIC-806, MA-13, IMC-47, NA-286, CAB-196, MA-14, SIC-23, SCA-5 for which random probabilities of identity were estimated below 1e-50 and only occasional mismatching SNPs were observed, usually involving a heterozygous vs. homozygous genotype. Unfortunately clones SIAL-70 and CCN-16 were not included in that study and no comparison was therefore possible to confirm the mislabeling.

The assessment of the correct germplasm identity of CEPEC’s collection is currently the focus of an ongoing project. Nevertheless, to be able to check the correct clonal identities, public databases of SNP profiles for trustworthy reference samples, using widely adopted SNP sets are necessary. An initial effort exists through the International Cocoa Germplasm Database at the University of Reading (www.icgd.rdg.ac.uk/). SNP data profiles for a few markers are downloadable. However, profiles for a much larger number of SNPs would be needed to allow effective data comparison. Concerted international efforts in this respect would represent an important advancement for the conservation, utilization and international exchange of cacao germplasm.

### Cacao clones display variable levels of SNP homozygosity

A wide distribution of observed homozygosity was seen in the sample of 90 clones with 16 clones showing less than 50% and 19 clones more than 90% of SNPs in a homozygous state (Fig 2). An overall highly significant inbreeding coefficient (F= 0.323 ± 0.002; p<0.001) was estimated, indicating an overall significant heterozygosity deficiency. Estimates of marker heterozygosity in cacao have been reported with RFLPs, microsatellites and SNPs in several studies in the last 25 years, usually showing heterozygote deficiency and significant inbreeding in several germplasm collections [38-42, 54-56]. Within the general theme of population analyses, DNA marker information has been used mainly to assess genetic diversity and structure of collections or natural populations or decipher the differential evolutionary origin and relationship of clones. Genome-wide heterozygosity estimates have been reported for a few clones [26] including TSH-1188 and CCN-51, also analyzed in our study. Specific estimates of microsatellite homozygosity showed that 20 out of 172 wild accessions from the Brazilian Amazon collection [57] and 163 clones out of 980 mainly wild or semi-domesticated accessions [58] had >90% homozygosity. No particular attention was usually given, however, to the potential application of individual plant marker data to inform deliberate inbreeding programs except for Efombagn et al. [40] who suggested that the more homozygous clones among the 400 surveyed should be used to generate uniform hybrids.

Besides the high efficiency for clone fingerprinting, the high polymorphism content of the SNPs used in this study, provides an effective tool to measure individual plant homozygosity. In other words, the ascertainment bias observed toward higher MAF SNPs (S3 Fig), should provide a slightly overestimated heterozygosity, or an underestimated, or more conservative, homozygosity. This contention is supported by considering the estimates of heterozygosity for clones TSH-1188 (0.343) and CCN-51 (0.431) reported earlier [26] based on transcriptome data. These values, converted to homozygosities, correspond to 0.657 and 0.569 respectively. Our homozygosity estimates from the 3,380 SNPs were considerably lower, estimated at 0.476 and 0.454 for TSH-1188 and CCN-51 respectively (Table 1). This comparison, albeit very limited, suggests that the estimates of homozygosity reported in our study might be underestimated and therefore on the conservative side. By extension, it is likely that all S_1_ and S_2_ plants analyzed could actually be even more homozygous than reported based on the 3,380 SNPs analyzed. Imputation of additional genome-wide SNPs using the available genome sequences could confirm this conjecture. In any case, more conservative estimates of homozygosity generated by this set of 3,380 SNPs would not compromise the objective of our proposed approach.

Despite the general conception that cacao is self-incompatible based on the favorable conditions for allogamy, the literature is rich on the fact that cacao may nonetheless present a high level of homozygosity even in wild and semi-domesticated populations. Up to 96% of self-fertilization in clones otherwise considered as self-incompatible, was reported [59]. This might be related to cacao’s distribution in its center of origin, frequently in populations isolated in river basins [60] and pollinated by insects (*Forcypomyia* spp) with small range of movement. Furthermore, semi-wild populations were established from small seed samples of the germplasm present in the source region [12], followed by occasional selection and expansion to the new areas. This is the case, for example, of the Amelonado population established in Bahia and in West Africa. Self-incompatibility is therefore not absolute. In fact, the existing variation for this trait has been the object of QTL mapping [61] and GWAS investigations [33, 34]. The observation of 55 out of the 90 clones with homozygosity above 70% in this study (S4 File) is therefore not at all surprising. Moreover, this result indicates that self-incompatibility and inbreeding depression might be less than a hurdle for the prospects of deliberately developing inbred lines in cacao.

### SNP data assisted hybrid breeding

Clones with high genome-wide homozygosity characterized in our study would be priority candidates to be assessed for general and specific combining abilities. They could be tested as parents of hybrids aiming to reduce variability in operational plantations and potentially exploit hybrid vigor in a more systematic way. A number of studies have shown a strong influence of dominance variance on cacao yield, supporting the dominance hypothesis for heterosis [15, 53, 62–65]. Additionally, a relationship between multivariate genetic divergence based on phenotypic traits with combining ability effects was reported in cacao, with the most divergent cultivar exhibiting high general combining ability, producing the best performing hybrids [66]. Crosses between cacao clones homozygous for alternative alleles at the largest proportion of SNPs, would maximize heterozygosity in the hybrid.

SNP heterozygosity in the hybrid could be used as a proxy to model heterosis for traits in which dominance variance is known to be important, as done in domestic animals [67–69]. The heatmap of genetic distances among the 90 clones and 49 inbred S_1_ and S_2_ individuals (S5 File), and more specifically between the most homozygous and divergent ones for alternatively fixed SNP alleles, could be used to propose experimental crosses to maximize heterozygosity in the F_1_ hybrid and test this hypothesis. Preliminary experimental evidence in support of such an approach in cacao indicated a significant positive correlation between genetic distance estimated with 96 SNPs and specific combining ability for yield [64]. The genetic distance estimator used in our study [47] attributes a four times higher weight to marker configurations of the type AA x BB or vice versa which result in 100% probability of heterozygosity in the F_1_ hybrid, when compared to configurations of the type AB x (AA or BB) or vice versa that only have 50% probability. Among the clones sampled in this study some are already widely used as parents of hybrids in Latin America. Given that breeders are frequently faced with an unfeasible large number of possible crosses, the data provided in this study could help establish some priorities.

### Homozygosity in inbred families

In all six inbred families studied, plants were observed with a nominal homozygosity higher than the expected mean, and in four of them these deviations were significant (Table 3). This is the first relevant observation to support our proposal of cacao inbred line development assisted by SNP markers. Even with a small number of individuals sampled, it is possible to find plants that deviate considerably from the mean family expected homozygosity. From the perspective of using SNP data to accelerate inbred cacao line development, it might be interesting to compare whether the within-family variation in attained homozygosity would be different in an S_1_ versus an S_2_ generation. One possibility is that more variation would be expected in the S_1_ generation as more genetic load would still be present for natural selection to operate. Unfortunately, we only sampled two S_1_ families, and the one derived by selfing clone SIAL-169 actually did not provide information, as the parental clone itself was already highly homozygous. Nevertheless, the range in the standardized proportions of homozygosity in the other S_1_ family (PS-13.19) was not different than in the four S_2_ families sampled (Table 2). This result suggests that the efficiency of using SNP data to select for individuals with higher homozygosity values should be similar at least in the first two generations of selfing. Evidently our study was limited not only to merely two generations, but also to a small number of parental clones to argue that this would be a general trend. In a follow up of this initial study, larger inbred families and more parental clones will be sampled.

A second important aspect of our data is revealed by the observed and expected homozygosities of the S_1_ individuals used to generate the S_2_ families. Both S_1_ individuals of clone TSH-1188 (families A and B) had a significantly lower than expected homozygosity, as indicated by the highly significant chi-squares (Table 1). On the other hand, the observed homozygosity of the S_1_ plant of CCN-51 was not different from the expected value. Important to note that the homozygosity of clones TSH-1188 (0.476) and CCN51 (0.454) are essentially equivalent (Table 3). These results indicate that the homozygosity of the S_1_ plants used considerably impacted the homozygosity levels achieved in the S_2_ families. While several S_2_ plants of CCN-51 reached a homozygosity above 0.870, the top homozygous S_2_ individuals of TSH-1188 only reached a value of 0.818. These results, illustrate the benefit of using SNP data to accelerate the production of cacao inbred lines. Had SNP data been available for a large S_1_ family of TSH-1188, the two S_1_ plants used to generate the S_2_ families probably would not have been selected to advance the program. Moreover, by selfing the top homozygous S_2_ plants of clone CCN-51, one should expect obtaining S_3_ plants with a homozygosity close or even above 95%. In other words, almost fully inbred lines would be produced with only three generations of selfing when assisted by SNP data. Evidently the larger the S_1_ and S_2_ families generated and genotyped, greater would be the opportunities to select outlier individual plants with higher proportions of homozygous SNPs.

### SNP assisted development of cacao inbred lines

Since the beginning of the adoption of cacao hybrid varieties, concerns were raised regarding the within-hybrid variability of hybrids produced from heterozygous parents [5, 6, 70]. Some methods were suggested to overcome this problem. First, based on the progeny segregation, more homogeneous clones were identified [13, 14, 71]. Some authors suggested observing the segregation in specific traits, and some partially inbred lines and their hybrids were tested [11, 14, 71], but variability persisted. Attempts were made to advance inbred lines, but the time, self-incompatibility and resource requirements could not be met by the underfunded programs, and were eventually discontinued [13, 72]. Alternative strategies included the search for haploid plants produced from “flat beans” [16]. Some haploid plants were found, diploidized and their hybrids resulted in high yielding and uniform progenies, while others presented poor performance [73]. Moreover, the occurrence of flat beans in cacao is very rare and the transmission of this trait to the offspring would be undesirable. A final strategy was the use of anther/ovule culture to produce double-haploid parents [17–19], however considerably more research and development will be required to make this technology operational.

Here we propose a simple and accessible alternative approach based on accelerating the development of inbred lines by regular selfing monitored by genome-wide SNP genotyping. This approach should be valuable for several other perennial outcrossing fruit and forest trees for which inbred line development has never been seriously considered. Initially, clones in a germplasm collection would be genotyped for a representative and large set of highly polymorphic SNPs and the standing homozygosity estimated. Clones that already enjoy high levels of homozygosity could be immediately included in a hybrid testing program and priority crosses among them selected based on estimating genetic divergence at SNP markers that would maximize heterozygosity in the F_1_ hybrid. Clones of major breeding interest but still showing considerable heterozygosity would be subject to self-pollination and tens of offspring produced. These offspring would be genotyped still at the seedling stage, parentage checked and the homozygosity estimated. Individual offspring with the largest departure from the mean family homozygosity would be top grafted to induce early flowering and prioritized to be further self-pollinated to advance the program. For example, in the S_1_ generation of PS-13.19, the low-end individuals had a homozygosity of 0.625 while the top ones 0.791. Assuming an equivalent standard deviation of homozygosity in the prospective S_2_ generation as in the S_1_, if the low-end individual is advanced, the expected homozygosity in the S_2_ would be 0.813, while if the high-end individual is used, a homozygosity of 0.896 is expected. However these would be only the average expected values. As pointed out above, transgressive homozygosity well above the expected mean could be found.

The proposal outlined above, evidently is simplistic and does not take into account all the key issues related to selection for phenotypic traits performance. Also, we have not assessed the potential relationships between individual SNP homozygosity and change in plant vigor due to inbreeding depression in the S_1_ and S_2_ generations. This will be the object of upcoming reports. Suffice it to say for now, that no substantial visual difference was seen in terms of overall plant vigor in the inbred individuals when compared to their parental clones. Reports on estimates of inbreeding depression following self-fertilization in cacao are scarce and limited to a few clones. The few reports have shown little if any inbreeding depression for a number of traits and occasionally even positive effects of inbreeding [65, 74, 75]. These observations, albeit requiring further validation, are in line with the fact that many currently planted clones already show moderate to high levels of inbreeding such as the Amelonado selection SIAL-169 (S4 File), resulting from differential inbreeding life histories. In fact, genome-wide data has shown that the domestication process of cacao has resulted in variable accumulation of deleterious mutations in different clones. While the Amelonado genome showed a distribution of deleterious mutations consistent with most of them having been purged by selfing, the Criollo populations have not undergone the same process [43]. Taken together, these data suggest that inbreeding depression will be variable across inbred families of different clones. Selection of inbred individuals to be advanced in each generation and crossed to generate hybrid trials will therefore involve not only maximum SNP homozygosity, but also selection for overall fitness, combination of key traits and adequate management of the inbred lines to be used in the production of commercial hybrid seeds.

The proposed approach of SNP assisted inbred line development in cacao presents several advantages. First, the cost of high-throughput DNA genotyping has dropped drastically in the last few years. The cost of genotyping a tree is considerably lower than the cost of blindly producing and advancing inbred individuals in the program by counting on a draw of luck in choosing the right inbred individual to advance. Second, highly homozygous seedlings can be selected in the nursery, reducing the number of plants taken to field trials. This has become particularly critical for species like cacao with long generation times, high demand of field areas and low level of mechanization. Several inbred seedlings can be top-grafted on well-cared flowering trees to speed up the buildup of inbreeding and reduce experimental area as suggested earlier [72]. With potential advancements in transgenic flower induction in cacao [76] trees carrying the FT (Flowering Locus T) gene could be used as rootstocks to induce even earlier flowering. Third, as shown by our results, even with only a few generations of selfing and a few offsprings, SNP data will indirectly allow exposing unfavorable alleles efficiently and these be eliminated by selection. Fourth, although this approach could be potentially carried out using microsatellite markers, it would be considerably less effective due to poor genome coverage. With the current prices of SNP genotyping, the cost advantage of microsatellite would be slim when compared to the large benefits of SNP data. By sampling a much larger portion of the genome, SNPs provide more extensive and accurate estimates of homozygosity, possibly also capturing polygenic effects for yield components. For improved homozygosity estimates, a further step in SNP genotyping beyond fixed-content arrays could consider low-pass whole genome sequencing combined with imputation for a much denser genome coverage [77, 78]. Considering that heterosis in cacao could potentially be modeled by the genetic divergence among lines and heterozygosity in the hybrid, the SNP genotyping data generated along the inbreeding process, could also be used to prioritize what cross combinations to test. Finally, the generation of hybrids from highly inbred lines not only should reduce unwanted variability in the farmer’s fields, but also in variety trials, improving the overall accuracy of selection in the breeding programs.

## Acknowledgments

We acknowledge the continued support for research provided by FAP-DF (Foundation for Scientific Research of the Federal District) through grants RECGENOMICS 00193-00000924/2021-92 and NEXTREE 0193.001.198/2016; FAPESB (Foundation for Scientific Research of the State of Bahia) through grant DTE0027/2013 and CNPq (Brazilian National Council for Scientific and Technological Development) productivity fellowships to KPG and DG. We would like to thank the field staff of CEPEC/CEPLAC for technical and logistic support for field sample collection.

## Supporting information

**S1 File.** Information for the Cacao SNPs. Correspondence between the EMBRAPA 65K Multispecies chip SNP codes EMB and the originally published SNPs Tcm id’s.

**S2 File.** SNP genotype data for 3,380 InfiniumII SNPs (A/G; A/C; T/G; T/C) for the 139 samples studied (90 clones and 49 S1 and S2 individuals). Data are presented in the Illumina AB format where the alleles A or T at the SNP correspond to the allele code “A” and alleles G or C at the SNP correspond to allele code “B”.

**S3 Fig.** Site frequency spectrum of the 3,380 SNPs in the 90 *Theobroma cacao* clones studied.

**S4 File.** SNP homozygosity estimates for the 90 cacao clones based on 3,380 SNPs.

**S5 File.** Genetic distance matrix. Matrix with overlapping heat map of the genetic distances among all 90 cacao clones and the 49 S1 and S2 offspring from clones TSH-1188, CCN-51, PS-13.19 and SIAL-169 based on 3,380 SNPs

**S6 File.** UPGMA dendrograms. (A) Dendrogram for the 90 clones only and (B) dendrogram for all 139 plants (90 clones and 49 inbred individuals) based on a matrix of genetic distances estimated with 3,380 SNPs.

## Data archiving

Genotype data are available in the supporting information S2 File.

